# Generation of an induced pluripotent stem cell line from a patient with immune checkpoint inhibitor-induced myocarditis and concurrent type I diabetes

**DOI:** 10.64898/2026.08.26.746482

**Authors:** Min Kyung Lee, Maria Rosaria Vitale, Yin Sun, Noah S. Wagner, Hiranya Amritavalli Sundar, Shuai Sun, Arav Ramchandran, Shaheen Khatua, Harrison Chou, Yushin Vivan Huang, Yan Zhuge, Joseph C. Wu, Han Zhu

## Abstract

Immune checkpoint inhibitor-induced myocarditis (ICIM) is a severe immune-related adverse event with heterogeneous clinical presentations and potential genetic susceptibility. Here, we established a human induced pluripotent stem cell (iPSC) line from an ICIM patient with an HLA-type distinct from previously reported line, who developed concurrent type I diabetes following ICI treatment. This line exhibited typical morphology, normal female karyotype, pluripotency, trilineage differentiation into all three germ layers, Sendai virus clearance, and no mycoplasma contamination. Given the fulminant nature and diverse clinical presentations of ICIM, expanding the repertoire of iPSC lines are critical for investigating ICIM heterogeneity and its underlying mechanisms.

**Resource Table:** 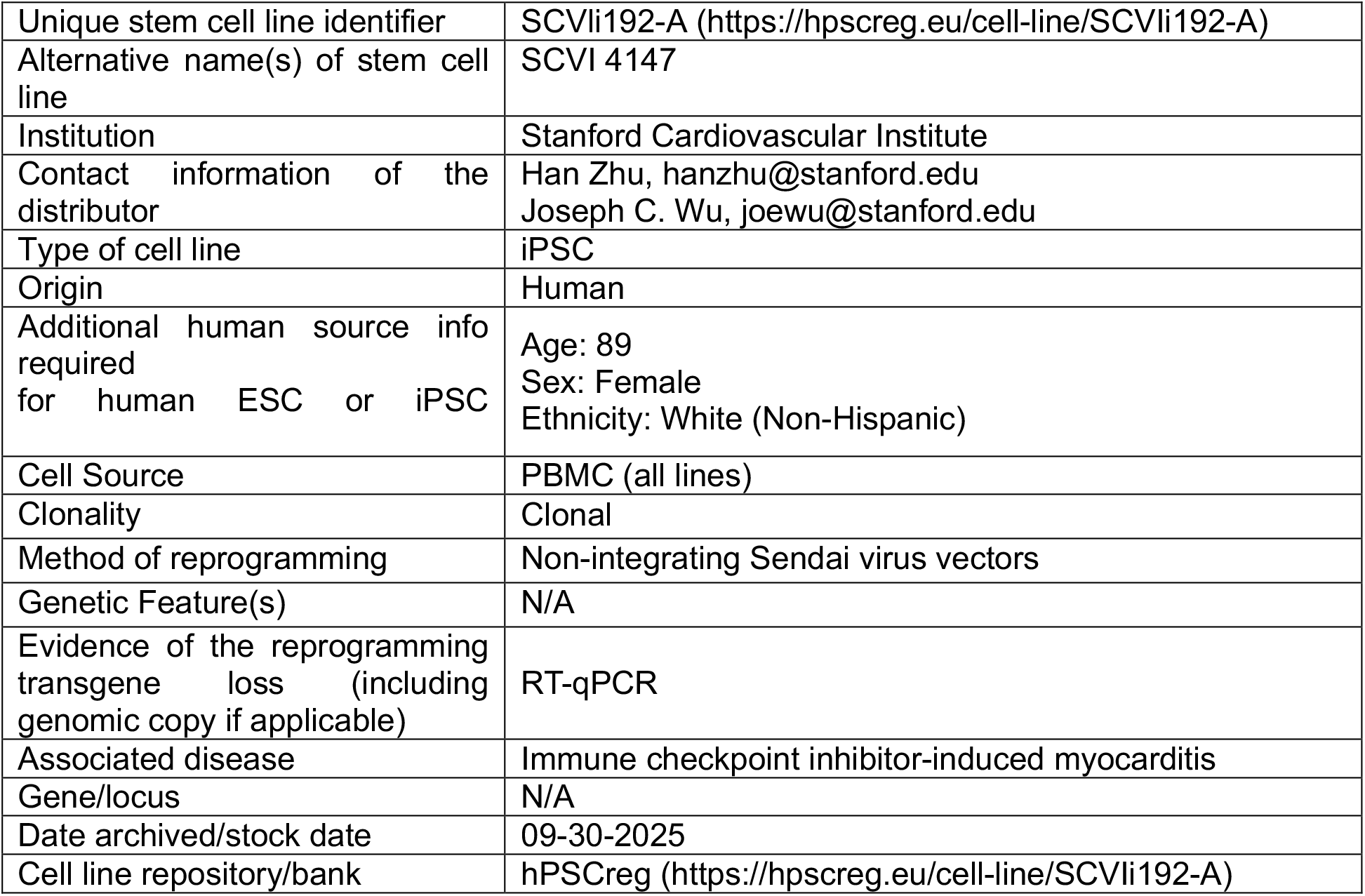

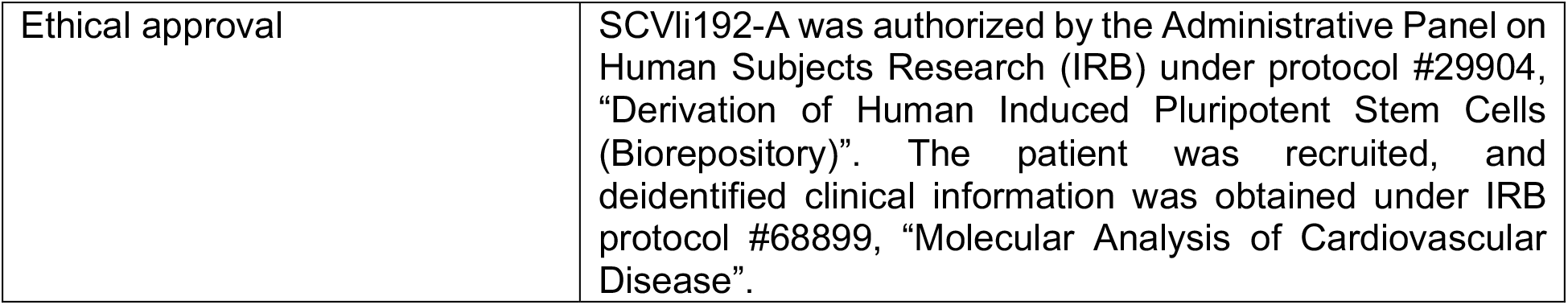

## 1. Resource Utility

This iPSC line was generated from a female ICIM patient with new-onset type I diabetes as a concurrent immune-related adverse event (irAE) following ICI therapy and has a distinct HLA type from the previously reported line. It will provide a resource for investigating clinical heterogeneity of ICIM and its underlying mechanisms.

## 2. Resource Details

Although immune checkpoint inhibitors (ICIs) have revolutionized the treatment of solid tumours by activating antitumor T cell immunity, they can cause immune checkpoint inhibitor-induced myocarditis (ICIM), a severe immune-related adverse event with reported case fatality rates of up to 40%^1-5^. Combination ICI therapy (anti-CTLA-4 plus anti-PD-1) is an established risk factor for ICIM^4,6^ while there are currently no established genetic risk factors that allow clinicians to identify patients at high risk prior to ICI therapy. However, emerging evidence suggests that patient-specific genetic factors, including human leukocyte antigen (HLA) type^7,8^ and clonal haematopoiesis of indeterminate potential (CHIP)-associated mutations^9^, may also contribute to disease susceptibility. Interestingly, patients with ICIM exhibit a broad spectrum of clinical presentations, including varying cardiac manifestations and concurrent immune-related adverse events^2-4^. These findings highlight the potential contribution of patient-specific genetic backgrounds to ICIM susceptibility and phenotypic heterogeneity, underscoring the need for experimental models that preserve interindividual differences among patients.

Patient-specific induced pluripotent stem cell (iPSC) lines provide a valuable platform for modeling ICIM *in vitro*, as they can be differentiated into various cardiac and immune cell types while retaining the donor’s genetic background^10^. Single cell multiomic studies of patients and preclinical mouse models have identified pathogenic T cell populations and complicate crosstalk among T cells, other immune cells, and cardiac cells that worsen cardiac inflammation^11-16^. Co-culture experiments using multiple iPSC-derived cell types, together with autologous HLA-matched T cells, could serve as an invaluable platform for investigating these multicellular interactions. Establishing a repository of iPSC lines representing diverse genetic backgrounds and clinical presentations could therefore advance our understanding of the mechanisms underlying ICIM heterogeneity, ultimately facilitating genetic driver-based risk stratification and the development of patient-specific therapeutic approaches.

Here, we established a human iPSC line (SCVli192-A) from an 89-year-old female ICIM patient who received Pembrolizumab as ICI treatment and develop concurrent type I diabetes. The iPSC line was generated by reprogramming the patient’s peripheral blood mononuclear cells (PBMCs) using a Sendai virus vector containing the four Yamanaka factors. The iPSC clone displayed typical morphology (Fig. 1A; scale bar: 530µM). The pluripotency markers, NANOG, SOX2, OCT3/4, were detected in the nuclei of the iPSCs by immunocytochemistry (Fig.1B; scale bar: 50µM). Reverse transcription quantitative polymerase chain reaction (RT-qPCR) showed markedly higher expression of NANOG and OCT3/4 in iPSCs than macrophages, further supporting their undifferentiated state (Fig. 1C). Sendai virus clearance was also confirmed in high-passage iPSCs by comparison with a validated early-passage iPSC line^17^ (Fig. 1D). Mycoplasma testing confirmed that the iPSC line was free of mycoplasma contamination (Fig. 1E). Short tandem repeat (STR) analysis proved the iPSC line had the same genetic origin with respect to the donor’s PBMCs. Furthermore, immunocytochemistry for ectoderm (PAX6 and OTX2), mesoderm (BRACHYURY and TBX6), and endoderm (FOXA2 and SOX17) markers demonstrated the capacity of this iPSC line differentiating into all three germ layers (Fig. 1F, scale bar: 100µM). Finally, G-banding karyotype assay revealed an apparently normal female karyotype (Fig. 1G).

**Fig 1.**
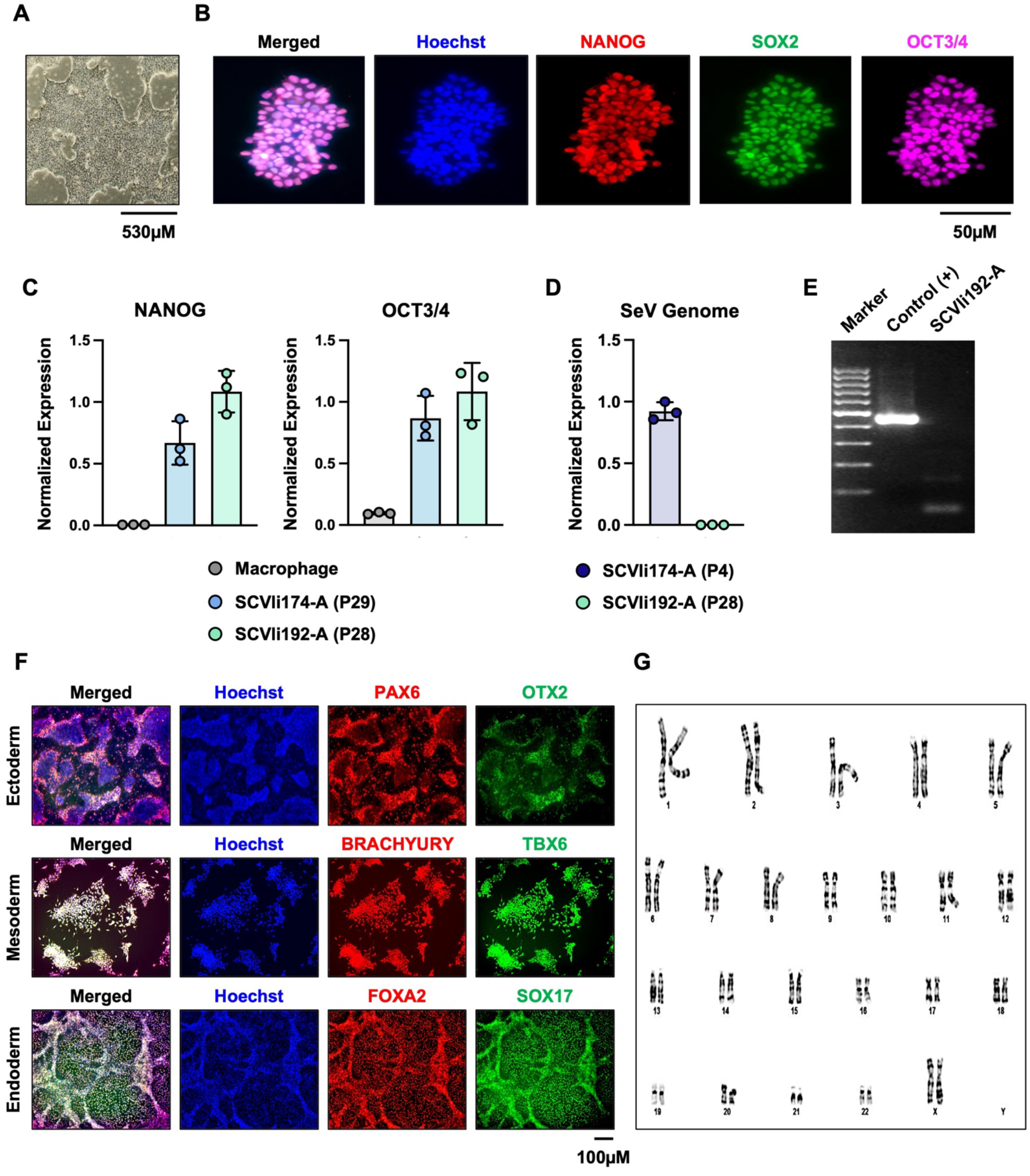
Characterization of an iPSC line, SCVli 192-A

## 3. Materials and Methods

### 3.1. Reprogramming

PBMCs were isolated from the whole blood of an ICIM patient using Percoll density gradient medium (GE Healthcare #17089109) and cultured in StemPro®-34 SFM medium (Thermo Fisher Scientific #10639011) with IL-6 (Thermo Fisher Scientific #PHC0063), IL-3 (PeproTech #200-03), and EPO (Thermo Scientific #PHC9631) (20ng/ml each) and FLT3 ligand (#PHC9414) and SCF (Peprotech #300-07) (100ng/ml each). PBMCs were reprogrammed using the CytoTune™-iPSC 2.0 Sendai Reprogramming Kit (Thermo Fisher Scientific #A16517) and plated on Matrigel (Corning #356231) in StemPro®-34 SFM medium. On day 7, the medium was switched to StemMACS™ iPS Brew XF medium (Miltenyi Biotec #130-104-368). Colonies emerging on days 10-15 were selected, expanded, and cryopreserved.

### 3.2. Cell Culture

All cell cultures were maintained at 37°C and 5% CO_2_ in a humified incubator. mTESR™ Plus medium (STEMCELL Technologies #05827) containing mTESR™ Plus 5x supplement (STEMCELL Technologies #05827) was used as the iPSC maintenance medium. Cells were cultured on Matrigel in iPSC maintenance medium until reaching 80% confluency. For passaging, cells were detached using 0.5mM EDTA and cultured in the presence of 10µM Rock inhibitor (MedChemExpress #HY10583). The medium was changed the day after passaging and every other day thereafter.

### 3.3. Immunofluorescent staining

Cells were fixed with 4% paraformaldehyde (Thermo Fisher Scientific #28908) for 15min, permeabilized with 0.1% Triton X-100 for 10min, and blocked with 1% BSA for 1hr. Primary and secondary antibodies (Table 2) were incubated overnight at 4 °C and room temperature, respectively. Nuclei were counterstained with Hoechst 33342 (Thermo Fisher Scientific #H3570), and images were acquired using a Keyence microscope.

**Table 1:** Characterization and validation.

| **Classification** | **Test** | **Result** | **Data** |
| --- | --- | --- | --- |
| **Morphology** | Photography Bright field | Visual record of the line: normal | Fig. 1 panel A |
| **Characterization of undifferentiated state** | Qualitative analysis: Immunofluorescence staining | Positive expression of pluripotency markers in the nuclei of iPSCs: NANOG, SOX2, and OCT3/4 | Fig. 1 panel B and C |
|  | Quantitative analysis: RT-qPCR | Positive expression of NANOG and OCT3/4 in the iPSC line but negative in differentiated macrophages |  |
| **Genomic stability** | Karyotype (G-banding), 1000X resolution | Normal karyotype: 46, XX | Fig. 1 panel G |
| **mtDNA analysis (IF APPLICABLE)** | Sanger sequencing, Deep sequencing, analysis software (Mitopore www.mitopore.de, mtDNA-Server 2, etc) | N/A | N/A |
| **Cell line identity verification** | STR analysis | 16 loci tested match well | Available with the authors |
|  | Microsatellite PCR (mPCR) OR SNP array | N/A | N/A |
| **Mutation analysis (IF APPLICABLE)** | Sequencing | N/A | N/A |
|  | Southern Blot OR WGS | N/A | N/A |
| **Microbiology and virology** | Mycoplasma | Luminescence: Negative | Fig. 1 panel E |
| **Characterization of the differentiated state** | Directed tri-lineage differentiation | Positive immunofluorescence (IF) staining of three germ layer markers | Fig. 1 panel F |
| **List of recommended differentiated state germ layer markers** | Expression of these markers has to be demonstrated at mRNA (RT PCR) or protein (IF) levels, at least 2 markers need to be shown per germ layer | Ectoderm: PAX6, OTX2<br>Mesoderm: BRACHYURY, TBX6<br>Endoderm: SOX17, FOXA2 | Fig. 1 panel F |
| **Donor screening (OPTIONAL)** | HIV 1 + 2 Hepatitis B, Hepatitis C | N/A | N/A |
| **Genotype additional info (OPTIONAL)** | Blood group genotyping | N/A | N/A |
|  | HLA tissue typing | N/A | N/A |

**Table 2:**
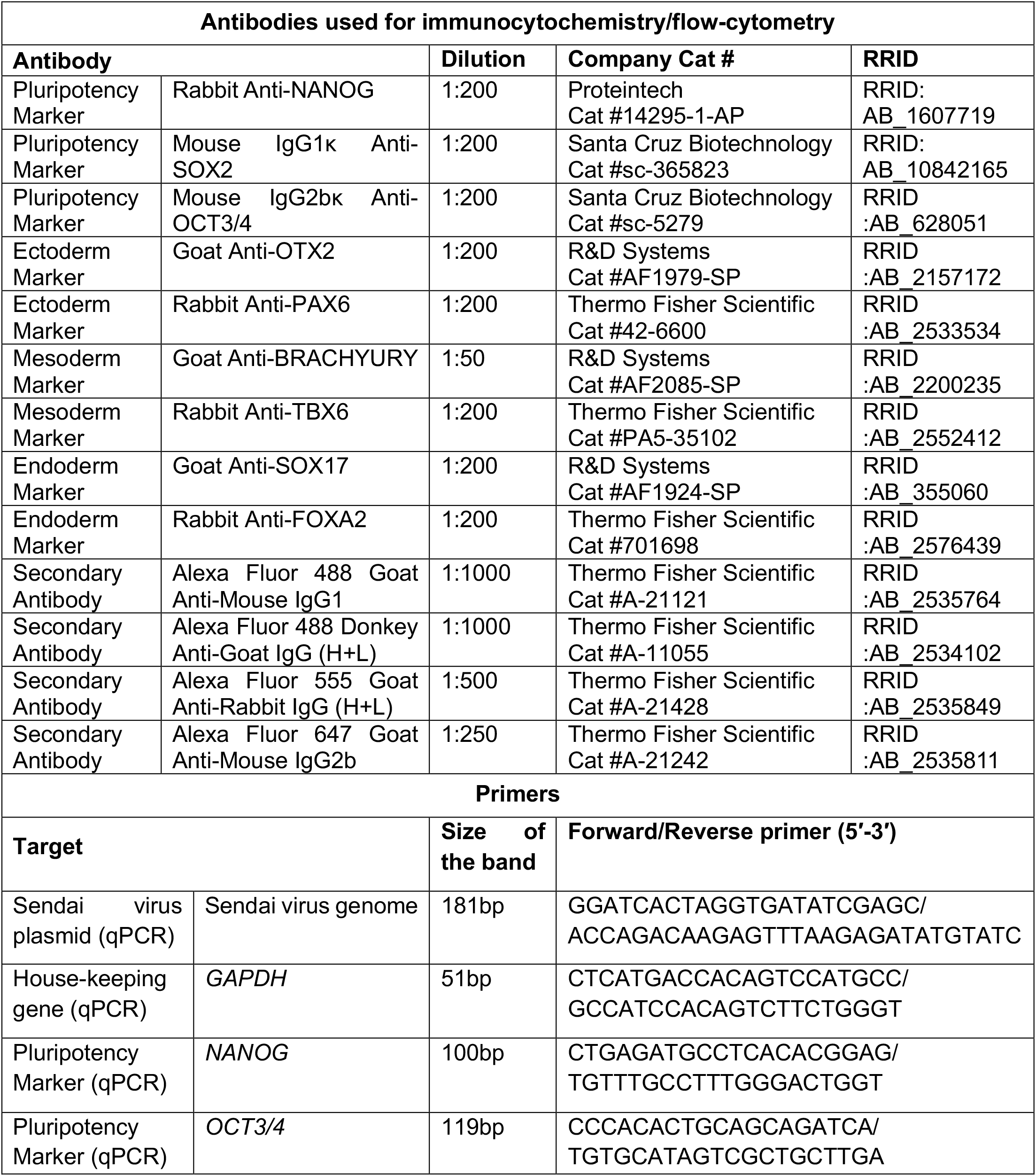
Reagent details.

### 3.4. RNA extraction and RT-qPCR

Total RNA was extracted using miRNeasy Micro Kit (Qiagen #217084). cDNA was synthesized using HiScript IV All-in-One Ultra RT SuperMix for qPCR (Vazyme #R433) and analyzed by qPCR using commercial primers (Table 2) and Taq Pro Universal SYBR qPCR Master Mix (Vazyme #Q712).

### 3.5. Trilineage differentiation

Ectoderm and endoderm differentiation was induced using the Stem MACS™ Trilineage Differentiation Kit (Miltenyi Biotec #130-115-660). For mesoderm differentiation, cells were cultured in iPSC maintenance medium containing 10 µM of Rock inhibitor for 24hr, followed by RPMI medium (Gibco #11875-093) with B27 minus insulin (Gibco #A18956-01) for 24hr and 6 µM CHIR99021 (Selleckchem #S2924) for an additional 24hr.

### 3.6. Short tandem repeat (STR) analysis

Genomic DNA was extracted from the iPSCs and donor PBMCs using the QIAamp DNA Micro Kit (Qiagen #56304). STR loci were amplified using the CLA IdentiFiler™ Plus PCR Amplification Kit (Thermo Fisher Scientific #A47624) and analyzed by capillary electrophoresis using an ABI 3730 Genetic Analyzer at the Stanford Protein and Nucleic Acid (PAN) Facility.

### 3.7. Karyotyping

Giemsa banding (G-banding) analysis was performed at Creative Bioarray. Cells were treated with 0.5µg/ml colcemid solution (Thermofisher #15210-040) for 3hr at 37 °C and 5% CO_2_, followed by hypotonic treatment with 0.075M KCl and fixation with methanol/acetic acid (3:1). Metaphase chromosome spreads were treated with 0.1% trypsin-EDTA and stained with Wright’s Giemsa stain. Chromosomes were analyzed using CytoVision karyotyping software (version 7.7, Leica Biosystems).

### 3.8. Mycoplasma detection

The iPSC line was tested for mycoplasma using the Mycoplasma PCR Detection Kit (abm #G238).

## 4. Discussion

ICIM is a severe cardiac complication of cancer immunotherapy, with case mortality rates approaching 40%. Furthermore, ICIM exhibits a broad spectrum of clinical presentations across patients, and growing evidence suggests that interindividual genetic variability may contribute this heterogeneity. However, no established genetic risk factors are currently available to identify patients at high risk of developing ICIM, limiting the safe administration of cancer immunotherapy. Establishing and expanding the human relevant models that preserve patient-specific genetic differences may therefore provide an invaluable resource for investigating the mechanisms underlying ICIM heterogeneity and ultimately contribute to the development of clinically applicable risk models.

Patient-derived iPSCs provide a valuable in vitro human model that preserves individual genetic backgrounds and can be differentiated into various cardiac and immune cell types. These cells can be further used in co-culture systems to investigate patient-specific pathogenic cellular interactions across multiple cell types. Importantly, given the central role of T cells in ICIM, co-culture with HLA-matched autologous T cells enables mechanistic investigation of patient-specific immune responses that affect T cell effector functions. Combined with advanced single-cell multiomic approaches that enable deep immunophenotyping, this platform can be a complementary human system for dissecting ICIM pathogenesis, for which mechanistic studies have relied largely on mouse models. Ultimately, this platform could facilitate the identification of novel therapeutic targets and large-scale screening of existing drugs, potentially enabling precision therapeutic strategies that account for the clinical and genetic heterogeneity of ICIM.

## CRediT authorship contribution statement

**Min Kyung Lee**: Writing-original draft, Visualization, Methodology, Formal analysis, Conceptualization. **Maria Rosaria Vitale**: Resources, Methodology, Conceptualization. **Yin Sun**: Resources, Methodology. **Noah S. Wagner**: Resources, Methodology. **Hiranya Amritavalli Sundar**: Resources, Methodology. **Shuai Sun**: Resources, Methodology. **Arav Ramchandran**: Methodology. **Shaheen Khatua**: Resources. **Harrison Chou**: Resources. **Yushin V Huang**: Resources. **Yan Zhuge**: Resources, Methodology. **Joseph C. Wu**: Resources, Funding acquisition. **Han Zhu**: Writing-review and editing, Supervision, Resources, Methodology, Funding acquisition, Data curation, Conceptualization.

## Acknowledgements

This study received funding from R01HL177581 (HZ), R01HL174432 (HZ), R01HL176822 (JCW), R03HL173146 (HZ), UM1TR006031 (JCW), K08HL161405 (HZ), Stanford CVI Seed (HZ), and AHA Transformational Project Award (25TPA1480322, HZ).

## Data availability

Data will be made available on request.

